# A 4D Bio-Kirigami Strategy for Engineering Complex Tissue Curvatures

**DOI:** 10.1101/2025.06.28.662122

**Authors:** Kaelyn L. Gasvoda, Aixiang Ding, Alexandria Sterenberg, Oju Jeon, Eben Alsberg

## Abstract

Four-dimensional (4D) systems offer a promising approach for generating sophisticated dynamic structures that mimic native tissue architectures. Among those, Kirigami strategies enable precise, localized control over morphing behaviors, yet their application in 4D tissue engineering remains unexplored. Here, we present a bio-Kirigami system that facilitates the creation of dynamic, transformable structures through spatially patterned hydrogels with distinct deformation modes dictated by encoded swelling differentials. The system consists of two biomaterial components: (1) a photocrosslinked hydrogel framework with controlled degradation and swelling behavior that drives the shape transformation and (2) a rigid support hydrogel frame. By leveraging photolithographic patterning, complex structures with continuously evolving configurations were achieved through preprogrammed deformations. Bio- Kirigami hydrogels encapsulating stem cells developed into tissue-like constructs with sophisticated configurations when cultured in tissue-specific environment. Notably, the engineered tissue constructs kept their shape integrity after excision from the outer support, demonstrating a robust platform for achieving intricate tissue curvatures.

## Introduction

Tissue curvatures[1, 2], arising from complex architectural arrangements[3], are prevalent across biological systems and span multiple length scales. These curvatures result from mechanical instabilities during tissue morphogenesis[4–6], including bending[2, 5], buckling[6], and creasing[7], and are intricately linked to tissue functionality. Mechanical cues such as stress and strain[8], generated by these curvatures[2], are transmitted through cell–extracellular matrix (ECM) interactions[9] and subsequently transduced into biochemical signals that regulate key biological processes[3], including cell alignment, migration, proliferation, and differentiation.

This dynamic and reciprocal interplay between mechanical forces[3, 4] and cellular responses helps orchestrate tissue organization and remodeling, wherein the mechanical cues influence cell behavior, and in turn, cells actively secrete, degrade, and/or remodel the ECM, establishing a feedback loop that profoundly influences both morphogenesis and pathophysiology[10].

Given their functional significance, recapitulating native dynamic tissue curvatures[11] and elucidating the biomechanical[2], biological[12], and biochemical[12] mechanisms governing their formation have become critical research priorities in developmental biology, offering valuable insights for tissue engineering[13]. However, current tissue engineering strategies predominantly employ cell-laden hydrogels in geometrically static configurations[14]. Although substantial efforts have been made to engineer hierarchical and composite hydrogel scaffolds that better mimic native tissue microenvironments[15], these approaches often fail to capture the dynamic architectural changes of developing and healing living tissues[16]. Therefore, integrating dynamic curvatures[13] into engineered tissues holds great potential for enhancing their physiological relevance and functionality.

Four-dimensional (4D) hydrogel systems have emerged as a promising strategy for engineering dynamically reconfigurable tissue architectures [17–19]. These systems typically consist of initially simple structures, such as strips, sheets, or discs, that undergo programmed shape transformations in response to external stimuli[20–23], forming complex architectures with continuous curvature variations. Incorporating live cells into these hydrogels and culturing them in tissue-specific environments enables the generation of dynamic curved tissues that are challenging to achieve using conventional or 3D bioprinting-based tissue engineering approaches. Recently, 4D Kirigami systems, which introduce strategically designed cuts into sheet-like structures, have been developed to enable out-of-plane transformations through localized and precisely controlled deformation[29–33]. Unlike conventional morphing hydrogels that exhibit limited deformation freedom due to continuous geometries and constrained interactions between neighboring regions, Kirigami-based architectures introduce region-specific morphing, enabling greater deformation amplitude and more complex, diverse configurations[32, 34]. This design strategy imparts programmable shape morphing along with superior mechanical properties, such as enhanced stretchability and reconfigurability, that may outperform traditional morphing hydrogels[35, 36].

Programmable Kirigami systems can generate tailored 3D architectures with large strains and enhanced conformability across multiple length scales[16, 32, 37, 38]. As a result, complex folded, bent, and twisted configurations can be readily achieved[39]. The high configurational versatility of 4D Kirigami systems allows for multiple degrees of deformation within a single construct. For instance, a Kirigami-engineered composite hydrogel membrane featuring laser-engraved periodic triangular cuts demonstrated deterministic, programmable out-of-plane deformation into 3D conical structures under uniaxial strain, enabling large, uniform, and reconfigurable shape changes across extended arrays[40]. In another example, Kirigami sheets integrating cuts with programmable folds achieved both compact and expanded 3D forms via folding-induced slit opening and reclosing, increasing the degrees of freedom for in-plane and out-of-plane actuation and enabling reconfigurable, stimuli-responsive transformations[41].

However, most current Kirigami systems are typically fabricated as thin films from materials such as metals[42], graphene[43], polymer composites[33, 38, 44], and elastomers [45, 46], using technologies like laser cutting, focused ion beam (FIB) writing, and 2D lithography. These materials and/or fabrication methods often suffer from limited cytocompatibility, restricting their capacity to support live-cell encapsulation. Consequently, applications have predominantly focused on photonics, electronics, and non-cellular biomedical devices like wearable sensors[47]. Their application for engineering dynamic tissue curvatures remains unexplored, primarily due to the stringent requirements for cytocompatibility and the need for continuous, controlled curvature evolution during tissue culture.

Here, we present a 4D bio-Kirigami system that integrates live cells to generate transformable, cell-laden architectures with dynamically evolving curvatures. This system consists of active cell-laden hydrogel components connected interiorly to an inert hydrogel ring (**Scheme 1**). The controlled swelling behavior of the active component, constrained by the surrounding ring, drives the formation of curved configurations. The bio-Kirigami system demonstrates dynamic shape transformations throughout the culture period due to hydrogel degradation and swelling, supports high cell viability, and facilitates tissue maturation, enabling the formation of structurally complex tissues with well-defined curvatures. Notably, following the excision of the deforming component, the engineered tissue constructs retain their shape for an additional week in culture. This bio-Kirigami system offers a robust platform for fabricating dynamic curved tissue constructs.

## Results

### Bio-Kirigami fabrication and characterization

The bio-Kirigami construct was fabricated using a multi-step photolithographic polymerization approach, as illustrated in Scheme S1 and Figure S1. The outer ring was composed of an interpenetrating network (IPN) hydrogel formed by calcium ion (Ca^2+^)-crosslinked alginate and photocrosslinked 8-arm polyethylene glycol (PEG) acrylate (ALG-PEG) [48], while the inner bars were fabricated using a composite hydrogel consisting of oxidized and methacrylated alginate (OMA) and gelatin methacrylate (GelMA) (OGMA). The chemical structures of 8-arm PEG acrylate, OMA, and GelMA were confirmed via **¹H NMR** analysis (Figure S2–S4).

Both ALG-PEG and OGMA hydrogels exhibited a higher storage modulus (G′) than loss modulus (G″) across a frequency sweep of 1–10 Hz (Figure 1a, b), confirming their solid-like mechanical characteristics. Notably, the ALG-PEG hydrogel displayed a higher G′ than OGMA, indicating greater stiffness. Over four weeks of culture in cell expansion medium, ALG-PEG maintained relatively stable swelling and degradation profiles, whereas OGMA underwent progressive changes in both properties (Figure 1c, d). In terms of swelling behavior, OGMA hydrogel exhibited a significantly higher swelling ratio compared to ALG-PEG (Figure 1d). Due to the differential swelling and degradation between these two hydrogel components, the inner bars were expected to undergo out-of-plane buckling deformation upon culture in media, leading to the formation of dynamically evolving 3D architectures.

**Figure 1.**
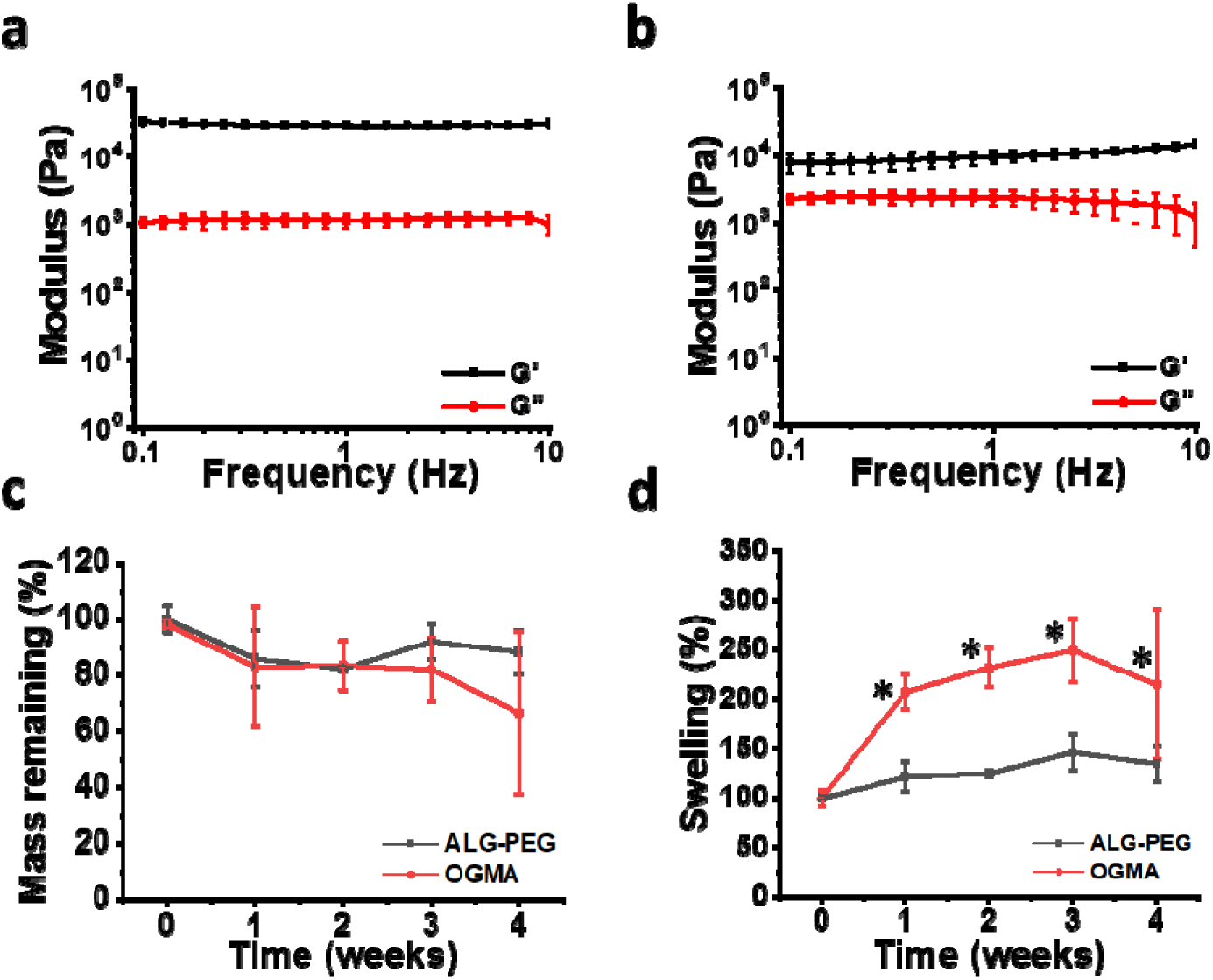
Rheological, degradation, and swelling properties. (a, b) Storage modulus (G′) and loss modulus (G″) of (a) ALG-PEG and (b) OGMA hydrogels as a function of frequency. (c) Mass degradation and (d) swelling of ALG-PEG (black) and OGMA (red) over 4 weeks of culture. *p < 0.05 when compared to other groups at the same timepoint. (a, b) N = 4. (c, d) N = 3.

### Shape morphing behavior of cell-laden bio-Kirigami constructs

Following the characterization of the polymer components within the bio-Kirigami system, its shape-transforming capability was then evaluated. The bio-Kirigami construct, laden with live NIH3T3 cells, was designed with a single inner bar connected to and surrounded by a ring-shaped outer frame (Figure S4). Non-cell-laden counterparts were included as controls. As anticipated, upon immersion in cell expansion media, the inner bars in both cell-laden and non-cell-laden systems exhibited out-of-plane buckling, forming arc-like deformations (Figure 2a, b). To quantify the dynamic shape transformation, the buckling height (z) was continuously monitored over 7 days. Both bio-Kirigami variants demonstrated a progressive increase in buckling height throughout the culture period (Figure 2c), driven by the constantly increasing swelling of the inner hydrogel components. Live/dead staining confirmed a predominance of live cells within the constructs, underscoring the cytocompatibility of the system for long-term culture (Figure 2d).

**Figure 2.**
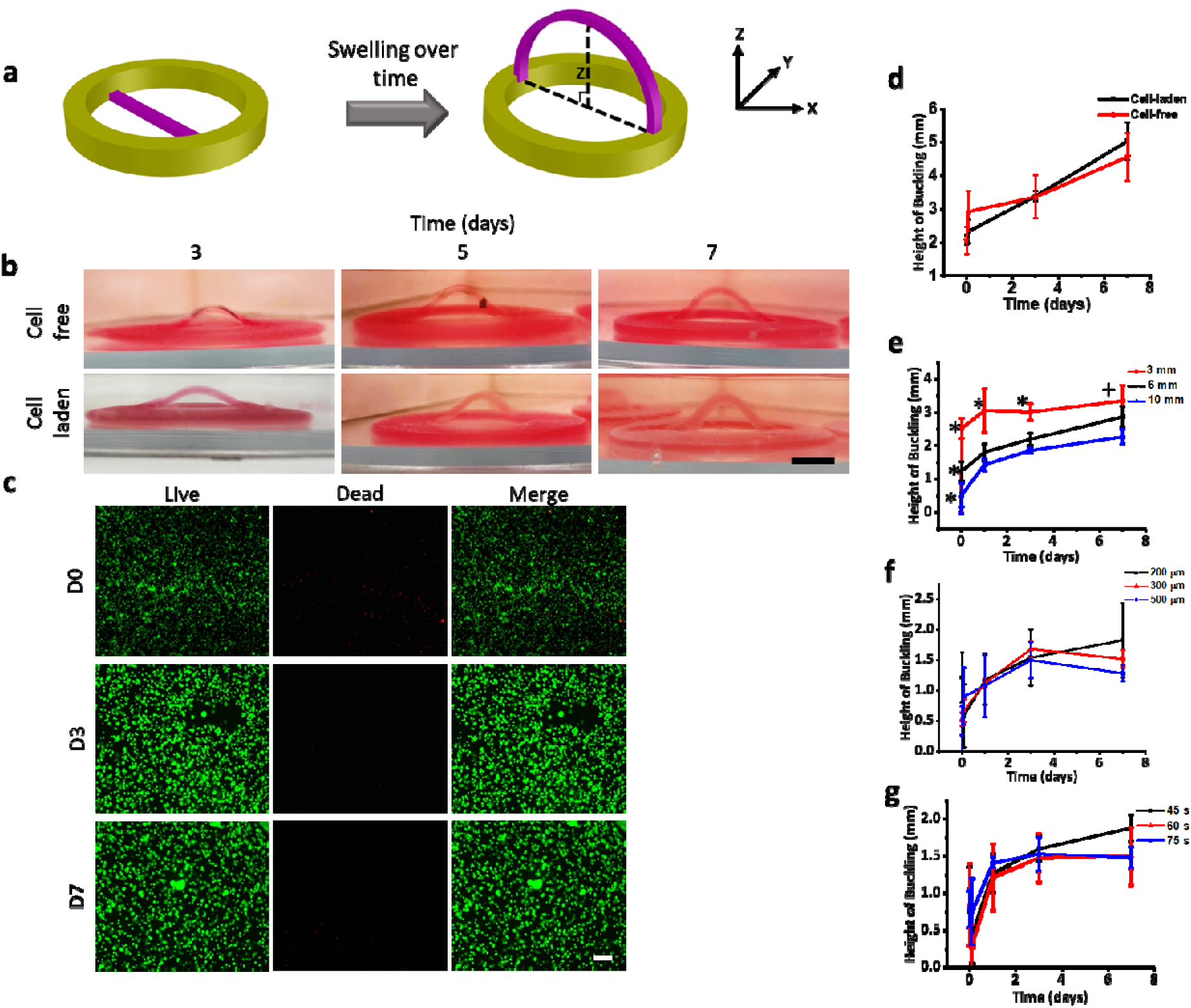
4D morphing properties of bio-Kirigami. (a) Illustration of morphing of bio- Kirigami containing a single strip during culture. (b) Representative images of non-cell-laden and cell-laden bio-Kirigami constructs at days 3, 5, and 7. NIH3T3 cells were encapsulated in the inner bar at a density of 5 × 10^6^ cells/mL hydrogel precursor solution. Inner bar dimensions: 15 mm × 3 mm × 0.3 mm. Outer support ring dimensions: Outer diameter: 20 mm; Inner diameter: 10 mm; Height: 0.7 mm. UV crosslinking time 45 s for inner bar and 5 min for outer ring at 20 mW/cm^2^. (c) Representative live/dead staining photomicrographs at days 0, 3, and 7. (d) Quantification of buckling height over culture time. (e–g) Influence of (d) inner bar width, (e) inner bar thickness, and (f) UV crosslinking time duration on the morphing behavior of bio- Kirigami constructs. *p < 0.05 compared to other conditions at the same time point. ^+^p < 0.05 compared to the 10 mm group at the same time point. Scale bars: (a) 5 mm; (c) 250 µm. N = 4.

The effects of key design parameters of the inner cell-laden bar on the 4D shape morphing behavior were further explored. Bar thickness (3 mm, 6 mm, and 10 mm), bar width (0.2 mm, 0.3 mm, and 0.5 mm), and ultraviolet (UV) crosslinking time (45 s, 60 s, and 90 s) of the constructs were varied, and buckling height was then measured over 7 days. Across all conditions, bio-Kirigami structures exhibited dynamic morphing behavior throughout the culture period. Increasing bar width (Figure 2e) resulted in a significant reduction in deformation, while variations in bar thickness (Figure 2f) and UV crosslinking time (Figure 2g) had no significant impact on the overall morphing capacity. Notably, a slight decrease in buckling height was observed at day 7 in constructs with bar thicknesses of 0.3 mm and 0.5 mm and in those crosslinked for 60 s and 75 s. This reduction in height is likely attributable to hydrogel degradation, leading to mechanical instability and subsequent structural relaxation.

### Intricate geometries formation

Upon confirming the feasibility of the approach, more intricate and distinctive geometric transformations, which were achieved through the strategic design of bio-Kirigami structures and programmed morphing, were explored. The initial geometries including “Cross,” “Double Strip,” “Single Dome,” “Concentric Double Dome,” and “Bolt” were fabricated using photolithography techniques. The dynamic evolution of these bio-Kirigami constructs throughout the culture period is depicted in Figures 3 and 4. All geometries, both cell-laden and non-cell-laden, exhibited continuous shape transformations throughout the culture period, consistent with our previous findings. The “Cross,” “Double Strip,” and “Single Dome” configurations exhibited single deformation patterns. Notably, the “Double Strip” construct exhibited a hierarchical morphing pattern, where one strip formed a larger buckling arc that overarched a second strip with a smaller buckling height. The “Bolt” configuration displayed a distinct non-uniform deformation pattern, in which the height of the swelling component was asymmetrically displaced to one side rather than remaining centered within the construct. By incorporating additional buckling structures into the inner component, such as in the “Spaceship” and the “Concentric Double Dome” designs, complex morphing patterns characterized by segmented curvature profiles were achieved.

**Figure 3.**
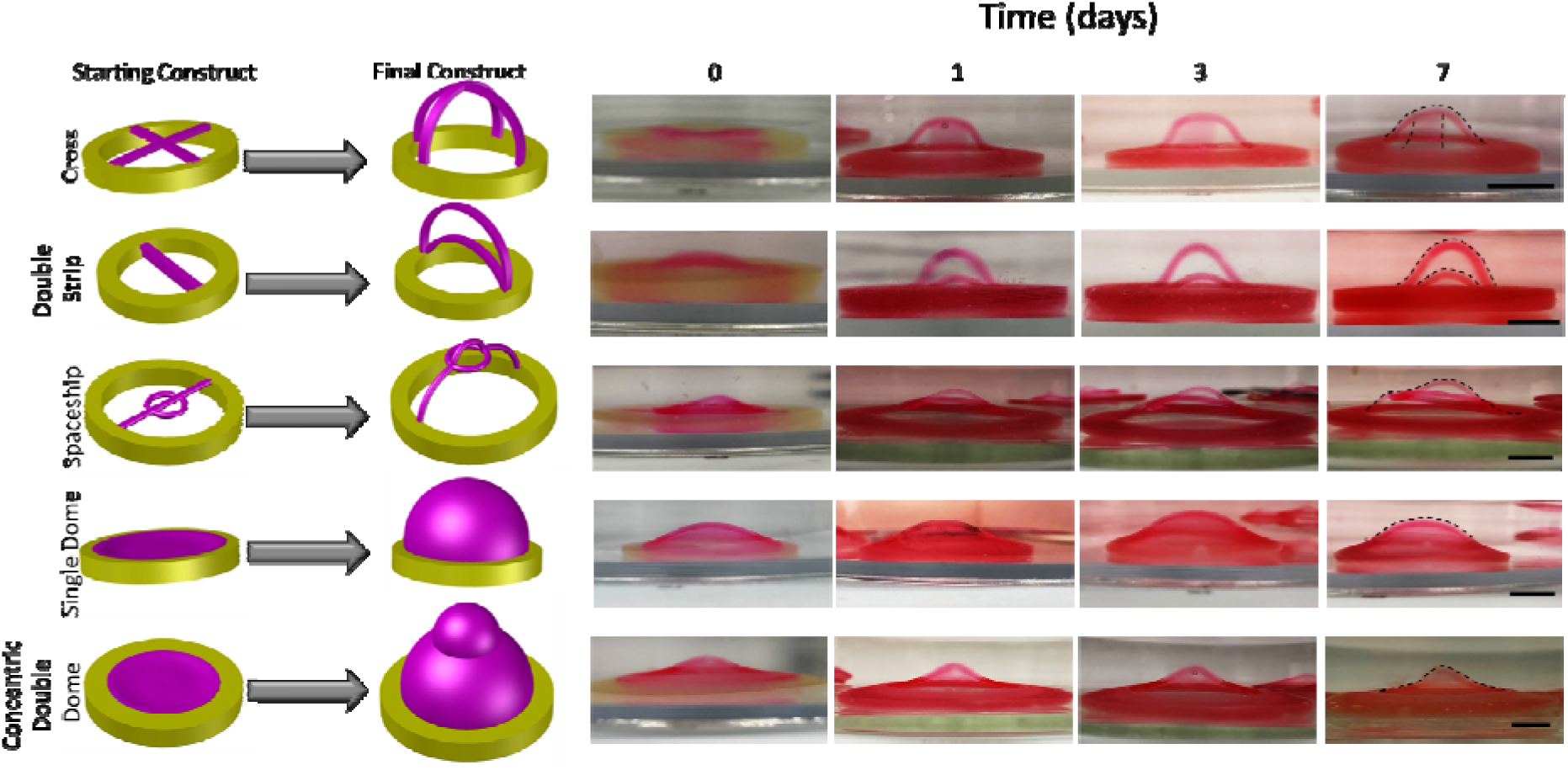
Complex configuration formation with non-cell-laden bio-Kirigami constructs. Representative Photographs at days 0, 1, 3, and 7 of culture in cell expansion media. Scale bars: 5 mm. N = 4.

**Figure 4.**
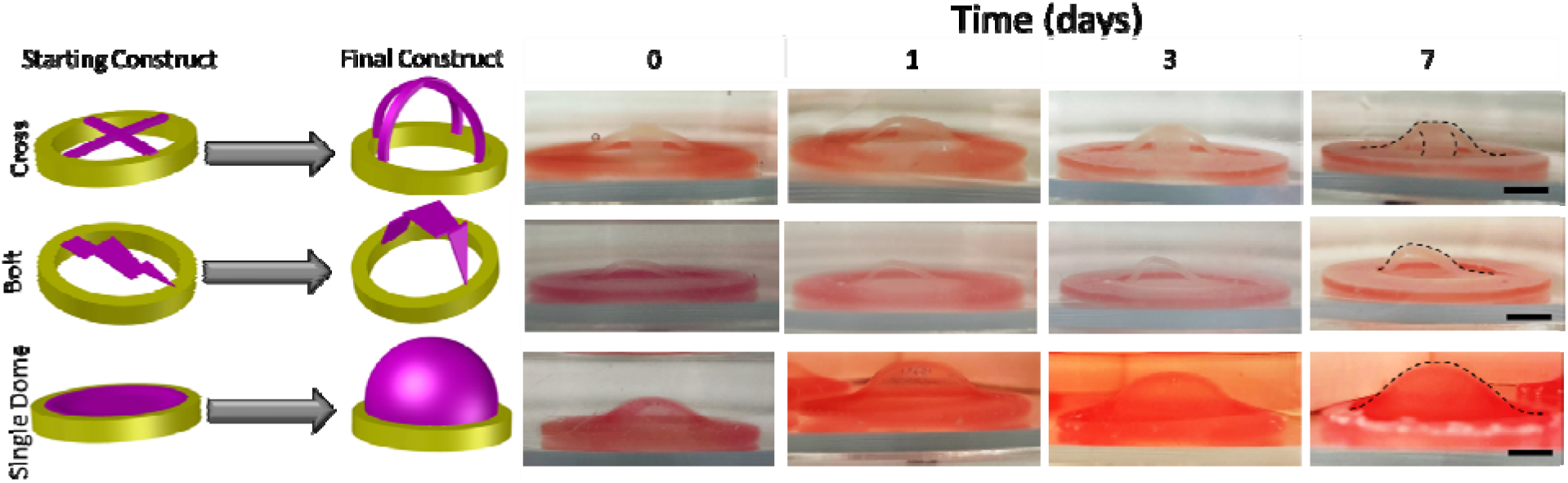
Complex configuration formation with cell-laden bio-Kirigami constructs. Representative photographs of complex constructs with encapsulated cells at days 0, 1, 3, and 7 of culture in cell expansion media. NIH 3T3 cells were encapsulated at a density of 10 × 10^6^ cells/mL OGMA polymer solution as part of the inner section (magenta). No cells were placed in the outer support ring (yellow). Scale bars: 5 mm. N = 4.

### bio-Kirigami-based 4D morphogenic tissue engineering

After completing the initial studies with encapsulated cells, the longevity of the bio-Kirigami system for 4D morphogenic tissue engineering applications was evaluated. Human mesenchymal stem cells (hMSCs) were encapsulated within the inner components of the “Single Dome” construct and divided into two groups: control constructs (Ctrl), which were cultured in standard cell-expansion medium, and differentiation constructs (CPM), which were cultured in chondrogenesis-pellet medium. Macroscopic images were captured at days 0, 7, 14, and 21 to monitor the shape evolution (Figure 5a). Throughout the culture period, both groups exhibited continuous morphing into dome-like configurations with progressively increasing deformation heights (Figure 5c). No significant differences in deformation height or morphing rates were observed between the two groups during this period.

**Figure 5.**
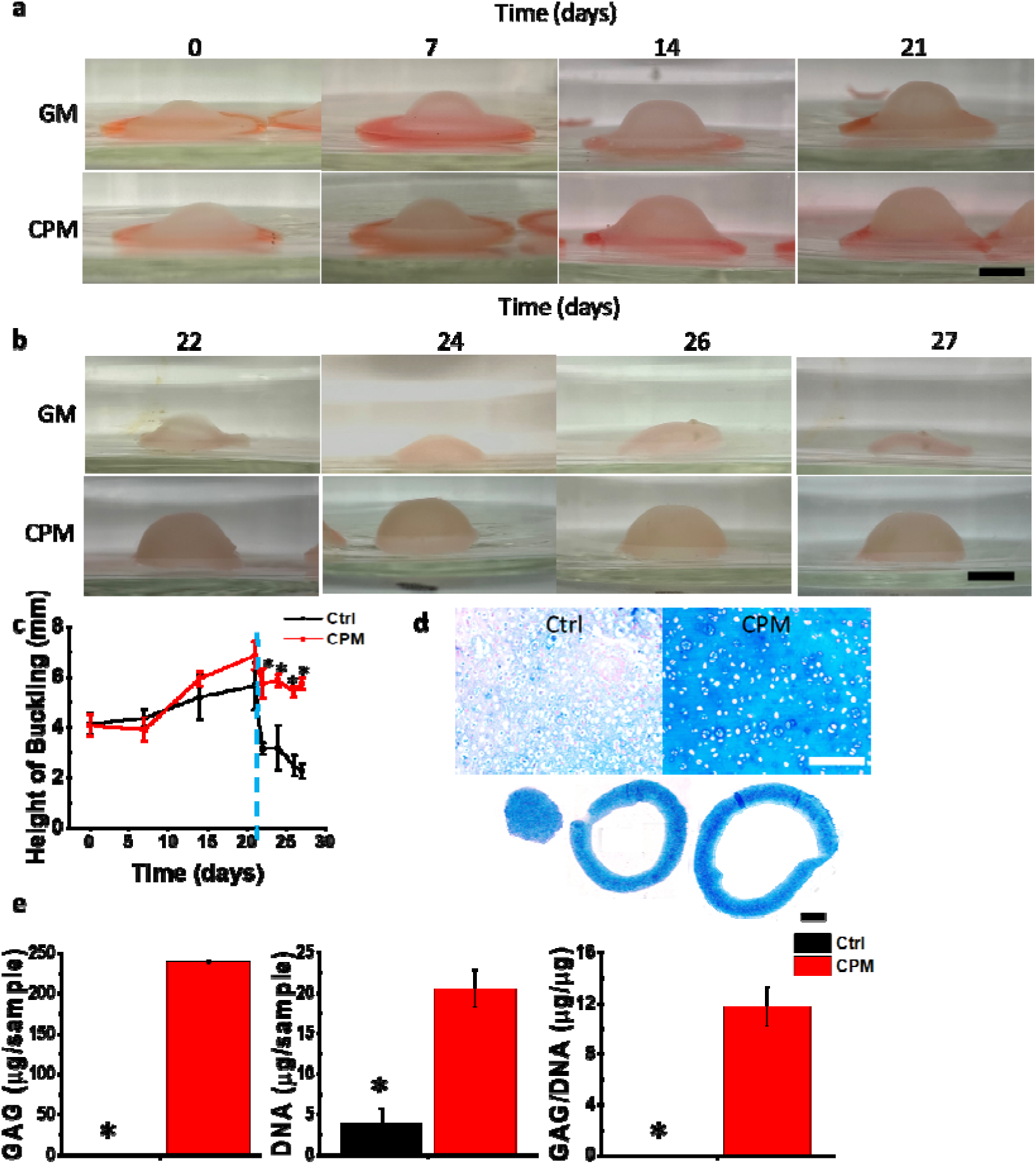
Chondrogenic differentiation of the 4D bio-Kirigami. (a) Representative macroscopic images of Ctrl and CPM constructs over 3 weeks of culture with the outer ring attached. Flexible component: diameter: 15 mm; height: 0.3 mm. Cell density: 40 × 10^6^ cells/mL hydrogel precursor solution. Crosslinked 45 s at 17 mW/cm^2^. Outer support ring: outer diameter: 15 mm; inner diameter: 8 mm; height: 0.7 mm. Crosslinked 5 min at 17 mW/cm^2^. Scale bar = 5 mm (b) Representative macroscopic images of Ctrl and CPM constructs over an additional 6 days of culture post-excision of the outer ring. Scale bar = 5 mm. (c) Quantitative measurements of deformation height. The blue dotted line indicated the time point when the outer support ring was removed, and the inner constructs continued to be cultured. (d) Upper panel: Alcian Blue (pH 0.2) staining of Ctrl and CPM constructs after 4 weeks of culture. Lower panel: Alcian Blue (pH 0.2) staining of CPM dome constructs at various cross sections of the differentiated dome. White scale = 250 µm. Black scale bar = 1 mm. (e) Biochemical analysis of DNA and GAG content in Ctrl and CPM constructs. Statistical analysis: p < 0.05. N = 2 for histology, N = 4 for other analyses.

Following the initial three-week culture, the deforming inner components were excised from the outer ring and cultured for an additional 6 days, during which time macroscopic images were acquired (Figure 5c). The results revealed that the Ctrl constructs gradually collapsed in the absence of the structural constraints of the outer ring, whereas the CPM constructs maintained their architecture and structural integrity, showing minimal visible and quantitative changes (Figure 5b, c). This stability in the CPM group was attributed to the substantial accumulation of ECM, which provided mechanical support and prevented structural collapse post-excision.

To further assess cartilage-like ECM formation, constructs at week 4 were subjected to histological and biochemical analysis. Alcian blue (pH 0.2) stain, which binds to sulfated glycosaminoglycans (GAGs) [49], a key biomarker of cartilage tissue, and not alginate, was performed to visualize chondrogenesis (Figure 5d). The CPM constructs exhibited intense staining, whereas the Ctrl constructs showed only weak staining (upper panel of Figure 5d). Additionally, strong Alcian Blue staining was observed throughout different locations within the CPM dome structure (lower panel of Figure 5d), suggesting the abundant GAG deposition and effective chondrogenic differentiation of the CPM constructs. This finding was further corroborated by biochemical quantification, which demonstrated significantly higher GAG content in the CPM constructs (Figure 5e). Interestingly, the CPM constructs also exhibited a significantly higher DNA content than the Ctrl constructs. This difference was attributed to hydrogel degradation in the Ctrl group, leading to structural collapse and cell leakage. In contrast, the CPM constructs, owing to their enhanced structural stability by cartilage-like ECM production, retained a greater number of cells, resulting in higher DNA content.

## Discussion

Previously, Kirigami strategies have been extensively explored in engineering fields in the design of metamaterials for optics[50] and wearable sensors[32]. The deformation of those Kirigami-based constructs is governed by the initial shape and “cutting” introduced during fabrication. Additionally, materials subjected to pre-stretching can undergo shape deformations upon strain release [30]. While Kirigami-inspired biomedical applications, such as silk fibroin-based scaffolds[35], have been reported, the integration of Kirigami strategies into hydrogel systems for engineering complex, dynamic tissues remains unexplored.

In this study, we introduce a 4D bio-Kirigami approach utilizing cytocompatible hydrogels to enable shape morphing without the need for pre-stretching or cutting. Instead, precise photolithographic patterning facilitates the fabrication of programmable structures with minimal processing steps, underscoring the simplicity and efficiency of the proposed system. To achieve preprogrammed shape transformations, two distinct hydrogel compositions with differential mechanical properties, swelling kinetics, and degradation behaviors (Figure 1) were employed. The first hydrogel, OGMA, exhibited rapid swelling in aqueous environments while maintaining sufficient mechanical strength to resist rupture. The second hydrogel, an ALG-PEG system [48, 51], demonstrated stable swelling and minimal degradation, providing structural reinforcement. The interplay between these two hydrogel systems allowed for controlled deformation while preserving overall structural integrity.

By leveraging these material properties and strategic structural designs, the bio-Kirigami constructs exhibited dynamic and continuous shape evolution over several weeks of culture (Figure 2). Variations in deformability were observed with changes in the width of the inner bar, whereas photocrosslinking duration and inner bar thickness did not result in statistically significant difference in deformability. Importantly, encapsulated cells remained highly viable and did not significantly alter the deformation compared to acellular constructs, highlighting the cytocompatibility of the system. This dynamic 4D bio-Kirigami platform thus offers a promising avenue for investigating the biomechanical influence of curvature on cellular behavior and tissue development.

The bio-Kirigami system supports both uniform and complex deformations across multiple degrees of freedom (Figure 3, 4). Importantly, intricate morphologies can be achieved through simple geometric design using single-step photolithography. Particularly notable is the system’s capacity to generate large, multiple curvatures with sophisticated patterns, independent of neighboring units, as demonstrated by the “Double Strips” and “Concentric Double Dome” designs. These configurations, which offer extensive deformation freedom, are challenging to achieve using other 4D strategies. The ability to program such intricate deformations underscores the versatility of the bio-Kirigami approach.

Beyond its capacity for structural transformation, the bio-Kirigami system facilitates dynamic 4D tissue engineering by supporting cell differentiation in a tissue-specific environment. Of particular significance, the deformed tissue constructs retained their structural integrity following excision from the supporting frame. For instance, in the “Single Dome” model (Figure 5), constructs cultured in chondrogenic medium initially relied on the outer ring for shape retention. However, upon removal of the ring, the matured tissue itself provided sufficient mechanical support to preserve the curvature. This ability to maintain post-excision structural integrity highlights the biomechanical stability and ECM-mediated reinforcement inherent in our system, positioning 4D bio-Kirigami as a powerful tool for engineering complex tissue architectures that are otherwise difficult to achieve using conventional methods.

Despite its advantages, the current system has several limitations that need further investigation. First, deformations in the fabricated constructs occur predominantly along the positive z-direction. However, applications in tissue morphogenesis and wound healing/tissue regeneration would benefit from bidirectional deformations along both positive and negative z-axes. This could potentially be achieved by predefining the swelling direction of the deforming components through multi-material layering or incorporating unit hinge designs [27]. Additionally, all reported constructs in this study utilized a circular outer support frame, yet alternative frame geometries could influence deformation patterns and expand the system’s capabilities. Future studies should explore the effects of different support frame designs on shape transformations.

Lastly, a persistent challenge in tissue engineering is the presence of residual hydrogel materials within the final constructs, which may interfere with cell-cell interactions and ECM deposition and remodeling while also posing potential immunotoxicity concerns[52]. Addressing this issue by mimimizing the amount of hydrogel component in the 4D bio-Kirigami constructs could enhance the translational potential of this approach.

To our knowledge, this 4D bio-Kirigami platform represents the first demonstration of a system capable of forming constrained curvatures, sustaining long-term cell viability, and supporting tissue regeneration. In addition to enabling the fabrication of complex tissue curvatures, such as dome-like structures resembling limb buds during embryonic development [53] or the skull cap and ends of long bone[54], this system has potential applications in tissue morphogenesis modeling due to its intrinsic dynamic nature. Furthermore, although not explicitly investigated in this study, the continuous shape evolution of these constructs could serve as a biomechanical cue for guiding tissue development, expanding their utility in regenerative medicine. Overall, the findings of this study establish 4D bio-Kirigami as a promising strategy for engineering dynamic, shape-morphing tissue constructs with high structural complexity and long-term stability. By further refining material compositions, design parameters, and scaffold degradation mechanisms, this platform holds great potential for advancing tissue engineering applications and regenerative medicine.

This study presents a 4D bio-Kirigami strategy that enables programmable, dynamic shape transformations for engineering complex tissue curvatures. By leveraging photolithographic patterning and distinct cytocompatible hydrogel compositions, bio-Kirigami constructs consisting of active inner morphing components fixed to an “inert” outer frame were fabricated, demonstrating controlled, continuous morphing without the need for pre-stretching or cutting. Through precise geometric design and the structural support of the outer frame, the system successfully generated complex tissue architectures with multiple degrees of curvature distribution. The 4D bio-Kirigami constructs supported long-term cell viability, differentiation, and tissue maturation while maintaining structural integrity post-excision, particularly highlighting the role of ECM-mediated reinforcement in sustaining morphological stability. This approach offers a promising avenue for advancing 4D tissue engineering by enabling the fabrication of complex architectures, mimicking morphogenesis, and providing geometric cues to guide tissue regeneration and beyond.

## Methods

### OMA Synthesis

OMA targeting theoretical 20% methacrylation and 5% oxidation was synthesized according to previously published protocols[55]. Briefly, 1% w/v sodium alginate (20 g, Protanal LF 20/40, FMC Biopolymer) was dissolved in deionized water overnight. Sodium periodate (0.218 g, Sigma, cat#311448-100G) was then added to this solution and left to react overnight in the dark. Next, 2-(N-morpholino)ethanesulfonic acid (19.52 g, MES, Sigma, cat#M8250-1KG) and sodium chloride (17.53 g, NaCl, Sigma, cat#S9888-25G) were added to the solution under vigorous stirring conditions. N-hydroxysuccinimide (1.77 g, NHS, VWR, cat#102614-812) and 1-ethyl-3-(3-dimethylaminopropyl)carbodiimide hydrochloride (5.84 g, EDC, Oakwood, cat#024810-250g) were added after the pH was adjusted to 6.5 using sodium hydroxide (10 M, NaOH, Sigma, cat#221465). After 10 min, 2-aminoethyl methacrylate hydrochloride (2.54 g, AEMA, PolySciences, cat#21002-10) was added to the solution and left to react overnight. The solution was then dialyzed using dialysis tubing of MWCO 3500 (Spectrum Laboratories Inc., cat#08-670-5B) for 7 days at 4 °C. The solution was then treated with activated charcoal (5 g, Neta Scientifics, cat#099536), frozen overnight at -80 °C, and lyophilized for 14 days (Labconco FreeZone Freeze Dry System, Kansas City, MO, model#7752020). ^1^H NMR confirmed the actual methacrylation was 7.63% (Figure S2).

### PEG-DA Synthesis

PEG-DA targeting 100% methacrylation was synthesized according to previously published protocols[56]. Briefly, poly(ethylene glycol) (10 g, PEG, Sigma, cat#202398-500G) was dissolved in toluene (90 mL, Sigma, cat#244511). Triethylamine (2 mL, Sigma, cat#T0886- 500ML) was then added to the solution as a catalyst. Acryloyl chloride (4 mL, Sigma, cat#A24109-100G) was added to the solution using a dropper while the solution was stirring vigorously on ice. After 15 min, the solution was placed in an oil bath (Kirkland, cat#1074184) at 45 °C to react overnight. The solution was then filtered and hexane (2 L, Sigma, cat#H306- sk4) was added under vigorous stirring conditions for 15 min. Next, the stirring was stopped, and the solution was left to settle for 30 min. The solvent was then removed. Ethyl ether (2 L, Sigma, cat#H306-sk4) was added to the precipitate under vigorous stirring conditions for 10 min. The solution was then left to settle without disturbance for 45 min. The solvent was removed, and the precipitate was left to air dry overnight. The precipitate was then rehydrated in diH_2_O (300 mL) and dialyzed using dialysis tubing of MWCO 3500 for 3 days at 4 °C. The polymer solution was filtered using a 0.22 µm pore filter membrane (Sigma, cat#S2GPT05RE) and frozen overnight at -80 °C. The frozen polymer was then lyophilized for 14 days. ^1^H NMR confirmed the actual methacrylation was 72% (Figure S3).

### GelMA Synthesis

GelMA targeting 100% methacrylation was synthesized according to previously published protocols[57]. Briefly, Gelatin (10 g, Type A, Sigma Aldrich) was dissolved in diH_2_O (100 mL) at 50 °C for 2 h. Next, methacrylic anhydride (10 mL, Sigma, cat#276685-100ML) was added dropwise to the solution and left to react at 50 °C for 1 h. The solution was left to cool naturally to room temperature and subsequently react overnight. The solution was then precipitated in excess acetone (4 L, Sigma, cat#179973-4L) and dialyzed using MWCO 12k-14k membrane tubing (Spectrum Laboratories Inc., cat#08-700-150) at 50 °C for 7 days. The polymer solution was then sterilized using a 0.22 µm pore filter membrane (Sigma, cat# S2GPT05RE) and frozen at –80 °C overnight. The frozen polymer was then lyophilized for 14 days to dry. The degree of substitution was determined by ^1^H NMR (Figure S4) to be 92%.

### Preparation of OGMA and ALG-PEG

The solvent for polymer dissolution was prepared in advance. Specifically, the photoinitiator 2- hydroxy-4’-(2-hydroxyethoxy)-2-methylpropiophenone (PI; 0.1% w/v, Sigma, cat# 410896-10g) was dissolved in Dulbecco’s Modified Eagle Medium with high glucose (DMEM-HG; Sigma, cat#D5523-50L) and stirred vigorously overnight in the dark. The resulting DMEM-HG/PI solution was sterilized using a 0.22 µm pore filter membrane and stored at 4 °C in the dark until use.

To prepare the OGMA polymer composite, synthesized OMA and GelMA were dissolved in DMEM-HG/PI at final concentrations of 4% w/v and 0.005% w/v, respectively. The solution was kept overnight in a 15 mL conical tube (Corning, cat#14-959-53A) to ensure complete dissolution.

For the ALG-PEG polymer composite, synthesized PEGDA was first dissolved in DMEM-HG/PI at 20% w/v overnight in a 15 mL conical tube. In a separate tube, unmodified alginate was dissolved in the same solvent at 1.25% w/v and similarly dissolved overnight. The next day, the PEGDA and alginate solutions were combined at a 1:1 volume ratio. Subsequently, 22.3 µL of a supersaturated calcium sulfate (CaSO_4_) solution (8.4 g in 40 mL of diH_2_O, Sigma, cat#AC315255000) was added to 1 mL of the PEGDA/alginate mixture. A dual-syringe mixing method was used to combine the solutions. Equal volumes of dissolved alginate and dissolved PEGDA were loaded into two 30 mL syringes (Fisher, Cat# 14-829-48), which were connected using a Luer-lock connector (Cole-Parmer, Cat#UX-50114-25). One syringe was fully depressed to transfer the solution into the opposing syringe, then reversed to return the solution to the original syringe. This back-and-forth mixing cycle was repeated 30 times. The solution was then allowed to rest in the dark for 10 minutes. This mixing–resting cycle was repeated three times.

The ALG-PEG solution was then ready for use.

### Degradation and Swelling Studies of ALG-PEG and OGMA Hydrogels

Degradation and swelling studies were conducted according to previously established protocols[58]. Briefly, ALG-PEG and OGMA precursor solutions were prepared as above were dispensed between two quartz plates separated by 1 mm spacers, forming circular disk geometries. The samples were photocrosslinked for 5 min under 20 mW/cm^2^ light intensity. After crosslinking, constructs were carefully removed from the quartz plates and transferred to 15 mL conical tubes containing 10 mL of DMEM-HG supplemented with 0.05% w/w sodium azide. At predetermined time points (Day 0, Week 1, Week 2, Week 3, and Week 4), constructs (N = 3 per time point) were collected and weighed to assess swelling. Samples were then frozen at -80 °C overnight and lyophilized for 3 days. The swelling (%) was calculated by the following equation:

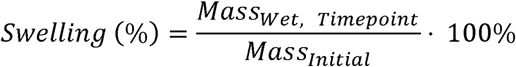

The degradation (%) was calculated by the following equation:

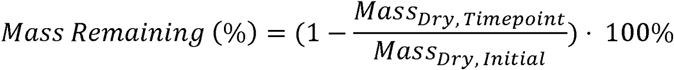

Rheological Characterization: Rheology studies were performed using previously established protocols[59]. Briefly, 6 mm diameter hydrogel disks with a height of 1 mm were fabricated and placed on a Kinexus rheometer stage with a 1 mm gap setting. Frequency sweep tests ranging from 0.1 to 10 Hz at 1% strain were performed at 25 °C to determine the storage (G′) and loss (G″) moduli of each hydrogel composition. *N =4*.

### Buckling Measurements of Deforming Component

As illustrated in Figure 2a, buckling measurements were performed by imaging the constructs submerged in 1X DPBS (300 mL; Sigma, cat#SH3001304) at a fixed distance of 5 cm using a phone camera. A calibration scale was also imaged at the same distance to define a known reference length (5 mm) in NIH FIJI software. This scale was applied to all construct images captured under identical conditions. For measurement, a horizontal baseline was drawn across the base of the deforming component to bisect the outer frame. A vertical line was then drawn from this baseline to the apex of the bent region within the construct. The length of this line, measured using the calibrated scale, was defined as the buckling height.

### Impact of Bar Thickness, Bar Width, and UV Crosslinking Time on Buckling

Rhodamine methacrylate (0.1% w/v; PolySciences, cat#669775-30-8) was added to the OGMA polymer composite for enhanced visualization. Constructs were fabricated to systematically investigate the effects of bar thickness, bar width, and UV crosslinking time on buckling behavior.

To assess the effect of bar thickness, the OGMA solution was sandwiched between two quartz plates using spacers of 0.2 mm, 0.3 mm, or 0.5 mm. A photomask with a rectangular cutout (15 mm × 3 mm) was placed on top, and the polymer was photocrosslinked for 45 s at 20 mW/cm^2^. The photomask and top quartz plate were removed, and the uncrosslinked polymer was washed away with 1× DPBS. Subsequently, ALG-PEG material was introduced around the rectangular OGMA bars, and a top quartz plate with 0.7 mm spacers was applied. A ring photomask (outer diameter: 20 mm; inner diameter: 10 mm) was positioned atop the assembly, followed by UV crosslinking for 5 min at 20 mW/cm^2^. The final construct was washed with 1X DPBS and transferred into a 6-well plate containing 8 mL of DMEM-HG supplemented with 0.05% w/w sodium azide. Constructs were imaged on days 0, 1, 3, and 7 using a phone camera (*N = 3*).

To investigate the influence of bar width, OGMA solution was dispensed between two quartz plates with 0.3 mm spacers. Rectangular photomasks with a length of 15 mm and varying widths of 3 mm, 6 mm, and 10 mm were used. The samples were photocrosslinked for 45 s at 20 mW/cm^2^. After removing the photomask and top quartz plate, uncrosslinked OGMA was washed away using 1X DPBS. ALG-PEG was then cast around the OGMA bars, and the top quartz plate with 0.7 mm spacers was positioned. A ring photomask (outer diameter: 20 mm; inner diameter: 10 mm) was applied, and samples were crosslinked for 5 min at 20 mW/cm². After washing with 1X DPBS, constructs were cultured in 8 mL of DMEM-HG containing 0.05% w/w sodium azide and imaged at days 0, 1, 3, and 7 using a phone camera (*N = 3*).

To determine the effect of UV exposure duration, the OGMA solution was placed between quartz plates with 0.3 mm spacers. A rectangular photomask (15 mm × 3 mm) was used, and the solution was exposed to UV light at 20 mW/cm^2^ for 45 s, 60 s, or 75 s. The photomask and top plate were removed, and uncrosslinked OGMA was washed with 1X DPBS. The ALG-PEG layer was cast around the OGMA bars and covered with a top quartz plate (0.7 mm spacers). A ring photomask was used for final crosslinking (5 min at 20 mW/cm^2^). Constructs were rinsed with 1X DPBS, transferred to 6-well plates with 8 mL of DMEM-HG containing 0.05% w/w sodium azide, and imaged at D0, D1, D3, and D7 a phone camera (*N = 3*).

### Cell Culture and Incorporation into Hydrogels

NIH 3T3 fibroblasts (ATCC, Manassas, VA) were expanded in Dulbecco’s Modified Eagle Medium-low glucose (DMEM-LG; Sigma, cat#D5523-50L) supplemented with fetal bovine serum (FBS; 10% v/v, Sigma, cat#18N103) and penicillin/streptomycin (P/S; 1% v/v, Gibco, cat#15140122). Cells were cultured at 37 °C in a humidified incubator with 5% CO_2_. Upon reaching 90% confluency, cells were harvested and suspended in the OGMA prepolymer solution at a density of 5 × 10^6^ cells/mL. The resulting cell-containing OGMA solution was then used to fabricate Kirigami constructs as described previously.

### Viability Staining

Cell viability within the constructs was assessed using a modified version of a previously established protocol[60]. Briefly, fluorescein diacetate (FDA; 1.5 mg/mL, Sigma, cat#F738-5G) and ethidium bromide (EB; 1 mg/mL, Fisher Scientific, cat#MP1ETBC1001) were combined by mixing 100 µL of FDA and 50 µL of EB with 30 µL of 1X PBS (pH 8) to prepare a working staining solution. A total of 150 µL of this solution was added to 5 mL of culture media containing a single construct. Constructs were incubated in the staining solution for 2 min at room temperature. Following staining, the media was removed, and the constructs were immediately imaged using a fluorescence microscope (Nikon Eclipse TE300, Japan). *N* = 3.

### Construction of Spaceship without Cells

OGMA solutions containing 0.1% w/v rhodamine methacrylate was dispensed between quartz plates separated by 0.3 mm spacers. A rectangular photomask (25 mm × 3 mm) was placed on top of the upper quartz plate, and the sample was photocrosslinked for 45 s under UV light at an intensity of 27 mW/cm^2^. Following crosslinking, the top quartz plate and photomask were removed, and the uncrosslinked OGMA was washed away using 1X DPBS. A second layer of OGMA was added around the crosslinked strip, and a new top quartz plate with 0.5 mm spacers was placed. A ring-shaped photomask (outer diameter: 15 mm; inner diameter: 10 mm) was aligned such that the central strip bisected the ring. The construct was then photocrosslinked at 27 mW/cm^2^ for 2 min. Uncrosslinked OGMA was washed away with 1X DPBS. Next, ALG- PEG solution was added around the OGMA construct, followed by placement of a quartz plate with 1 mm spacers. A larger ring-shaped photomask (outer diameter: 30 mm; inner diameter: 20 mm) was applied, and the construct was crosslinked under 27 mW/cm^2^ UV light for 5 min. The remaining uncrosslinked polymer was washed away with 1X DPBS. Constructs were cultured in 6-well culture plates containing 8 mL of DMEM-HG supplemented with 0.05% w/w sodium azide. Macroscopic images were taken using a phone camera on days 0, 1, 3, and 7 (*N = 4*).

### Construction of Concentric Double Dome with Cells

OGMA solutions containing 0.1% w/v rhodamine methacrylate were loaded between quartz plates separated by 0.4 mm spacers. A circular photomask (diameter: 8 mm, Figure S5) was placed on top, and the solution was photocrosslinked for 60 s at 22 mW/cm^2^. The photomask was then replaced with a ring-shaped mask (outer diameter: 25 mm; inner diameter: 5 mm, Figure S5), ensuring that the previously crosslinked circular region was centered within the ring. This secondary region was crosslinked for 180 s at 22 mW/cm^2^. Uncrosslinked OGMA was washed away with 1X DPBS. ALG-PEG solution was then placed around the OGMA hydrogel, and a top quartz plate with 1 mm spacers was applied. A final ring-shaped photomask (outer diameter: 30 mm; inner diameter: 20 mm, Figure S5) was used to crosslink the material for 5 min at 22 mW/cm^2^. Uncrosslinked material was washed away with 1X DPBS. Constructs were cultured in 6-well culture plates with 8 mL of DMEM-HG containing 0.05% w/w sodium azide. Imaging was performed macroscopically on D0, D1, D3, and D7 using a phone camera (*N = 4*).

### Construction of Double Strip without Cells

OGMA solutions containing 0.1% w/v rhodamine methacrylate were added between quartz plates with 0.3 mm spacers. A rectangular photomask (15 mm × 3 mm, Figure S5) was applied, and the solution was crosslinked for 45 s at 21 mW/cm^2^. After rinsing with 1X DPBS, the first strip was set aside. A second OGMA strip of identical dimensions was then crosslinked for 90 s under the same conditions and similarly washed. The two OGMA strips were stacked, with the longer-exposed (stiffer) strip placed at the bottom. ALG-PEG material was added to encapsulate the assembly, and a quartz plate with 1.7 mm spacers was placed on top. A ring-shaped photomask (outer diameter: 20 mm; inner diameter: 10 mm, Figure S5) was applied, and the sample was crosslinked for 5 min at 21 mW/cm^2^. Following washing with 1X DPBS, the constructs were cultured in 6-well culture plates with 8 mL of DMEM-HG medium containing 0.05% w/w sodium azide. Macroscopic images were taken at D0, D1, D3, and D7 using a phone camera (*N = 4*).

### Construction of Complex Deformations with Cells (Cross, Single Dome, Bolt)

NIH 3T3 cells were suspended in the OGMA solution at a final density of 10 × 10^6^ cells/mL. The cell-laden OGMA solutions were then loaded between quartz plates separated by 0.3 mm spacers. Photomasks corresponding to three geometries, including Cross (3 mm × 15 mm, Figure S6), Single Dome (diameter: 15 mm, Figure S6), and Bolt (15.75 mm × 6.60 mm; designed using PowerPoint’s default shape tools, Figure S6), were positioned atop the quartz plate and exposed to UV light for 45 s at an intensity of 22 mW/cm^2^ to induce photocrosslinking. Following crosslinking, the top plate and photomask were removed, and uncrosslinked OGMA was rinsed away with 1X DPBS. ALG-PEG precursor solution was placed around the OGMA structures, and a top quartz plate was placed on top of the prefabricated construct with 0.7 mm spacers. A ring-shaped photomask (outer diameter: 20 mm; inner diameter: 10 mm, Figure S6) was positioned on the top quartz plate to center the OGMA hydrogel within the ring, and the ALG- PEG solution was photocrosslinked for 5 min at 22 mW/cm^2^. The constructs were cultured using the conditions described above. Constructs were imaged at days 0, 1, 3, and 7 using a phone camera (*N = 4*).

### Chondrogenic Kirigami Constructs

hMSCs were expanded in DMEM-LG supplemented with 10% v/v FBS, 1% v/v P/S, and 10 ng/mL fibroblast growth factor-2 (FGF-2, R&D, cat#233-FB-MTO) in a humidified incubator at 37 °C with 5% CO_2_, with medium changes every other day. Upon reaching 80% confluency, cells at passage 4 were harvested for encapsulation. Cells were encapsulated within the OGMA polymer solution at a density of 40 × 10^6^ cells/mL. The cell-laden solution was loaded between quartz plates separated by 0.3 mm spacers. A circular photomask (diameter: 10 mm, Figure S7) was applied and the construct was crosslinked for 45 s under UV light at 17 mW/cm^2^.

Uncrosslinked polymer was removed by washing with 1X DPBS. ALG-PEG precursor solution was subsequently added around the crosslinked OGMA construct, and a top quartz plate was placed with 0.7 mm spacers. A ring-shaped photomask (inner diameter: 8 mm; outer diameter: 15 mm, Figure S5) was used to define the surrounding ALG-PEG frame, and the OGMA was crosslinked under UV light for 5 min at 17 mW/cm^2^. Unreacted material was removed by washing with 1X DPBS.

Constructs were cultured in either expansion medium or chondrogenic differentiation medium, based on previously established protocols[49]. The expansion medium consisted of DMEM-LG supplemented with 10% v/v prescreened FBS and 1% v/v P/S, while the differentiation medium consisted of DMEM-HG supplemented with 1% v/v ITS+ Premix (Fisher, cat#CB40352), 1% v/v P/S, 100 mM sodium pyruvate (Fisher, cat#SH3023901), 1% v/v non-essential amino acids (Gibco, cat#11140050), 37.5 µg/mL L-ascorbic acid (Wako USA, cat#013-12061), 100 nM dexamethasone (Sigma, cat#D4902-100mg), and 10 ng/mL transforming growth factor-β1 (TGF- β1, Peprotech, cat#100-21). Cell encapsulation was performed using OGMA solution at a density of 40 × 10^6^ cells/mL to form Kirigami constructs.

Both culture media (6 mL) were changed every other day. Constructs were macroscopically imaged using a phone camera at days 0, 7, 14, and 21. After 3 weeks, the constructs were excised from the surrounding ALG-PEG ring and cultured for an additional week. During this final week, macroscopic images were acquired at days 22, 24, 26, and 27. At the end of the culture period, constructs were collected for biochemical assays (*N = 4*) and histological analyses (*N = 2*).

### Bending Angle Calculation for Excised Deformed Constructs

Bending height measurements were performed using the same method as applied to non-excised constructs. Each construct was placed on a flat culture dish containing 1X DPBS (300 mL) and imaged macroscopically from a fixed vertical distance of 5 cm. A calibration scale was imaged under identical conditions to ensure consistent spatial referencing. Constructs were returned to the culture medium after imaging for further culture. For image analysis, the known length of the calibration scale was first measured using the line tool in NIH FIJI (ImageJ) to establish a pixel-to-length conversion factor. Buckling (bending) height for each construct was then measured using the same tool and calibration parameters. *N = 4*.

### Biochemical Analysis

DNA and GAG contents were analyzed using previously reported protocols[61]. Briefly, the samples were digested overnight at 65 °C in a digestion solution (pH 6.5) consisting of papain (25 µg/mL, Sigma, cat#P4762), L-cysteine (2 mM, Sigma, cat#C7352-25g), sodium phosphate(Na_2_HPO_4_, 50 mM, Fisher, cat#BP332-500), Ethylenediaminetetraacetic acid disodium salt dihydrate (EDTA, 2 mM, Fisher, cat# S25687). DNA was determined by measuring fluorescence intensity at an excitation of 488 nm and emission 520 nm on a plate reader (iD5, Molecular Devices, San Jose, CA) using a Quant-iT PicoGreen dsDNA reagent (Luminoprobe, cat#42010) and a 20X TE buffer, Rnase-free (Life Technologies, cat# T11493). GAG was determined by measuring the absorbance of the samples at 595 nm after reacting the solution with 1,9-dimethylmethylene blue (DMMB, Sigma, cat#341088-1G) dye on a plate reader. *N = 4*.

### Histological Analysis

Constructs were collected after 27 days of culture and processed for paraffin embedding and sectioning following previously established protocols[62]. Briefly, samples were fixed in 4% w/w paraformaldehyde (Sigma, cat#P6148-500G) overnight at room temperature. Fixed tissues were dehydrated through a graded ethanol series with sequential 2 h incubations in 70%, 95%, and 100% v/v ethanol (EtOH, Fisher, cat#SF1C163). Following dehydration, samples were incubated in a 1:1 mixture of ethanol and xylene (Fisher, cat#X3S-4) for 2 h, followed by two additional 2 h incubations in 100% xylene. Samples were embedded in paraffin (Epredia, cat#8336) and sectioned at 7 µm thickness. Sections were stained for GAG content using Alcian Blue (pH 0.2, Fisher, cat#92-31-9) and counterstained with Nuclear Fast Red (Polysciences, cat#09773) according to published protocols[63]. Alcian Blue was applied for 30 min, followed by a 5 min incubation with Nuclear Fast Red. Stained sections were subsequently mounted and imaged using a light microscope (Nikon Eclipse TE300, Japan) equipped with a14MP Aptina Color CMOS digital camera (AmScope, Irvine, CA). *N = 2*.

### Statistical Analysis

All data are reported as mean ± standard deviation (±SD). Significance was determined using one-way ANOVA with a post-hoc Tukey HSD test. p < 0.05 was considered significant unless otherwise specified.

## Author contributions

**K.L.G.:** Methodology, Investigation, Formal Analysis, Visualization, Writing-Original Draft, Writing – Review & Editing. **A.D.:** Conceptualization, Methodology, Formal Analysis, Writing-Original Draft, Writing – Review & Editing. **A.S.:** Investigation. **O.J.:** Methodology. **E.A.**: Conceptualization, Methodology, Formal Analysis, Resources, Supervision, Funding acquisition, Writing – Review & Editing.

## Acknowledgements

The authors gratefully acknowledge funding support from the Department of Veterans Affairs, Veterans Health Administration, Office of Research and Development, Rehabilitation Research and Development Service under award numbers RX004288 and RX004825 and the National Institutes of Health’s National Institute of Arthritis and Musculoskeletal and Skin Diseases under award number R01AR081448. The contents of this publication are solely the responsibility of the authors and do not necessarily represent the official views of the Department of Veterans Affairs or the National Institutes of Health.

---

**Scheme 1.**
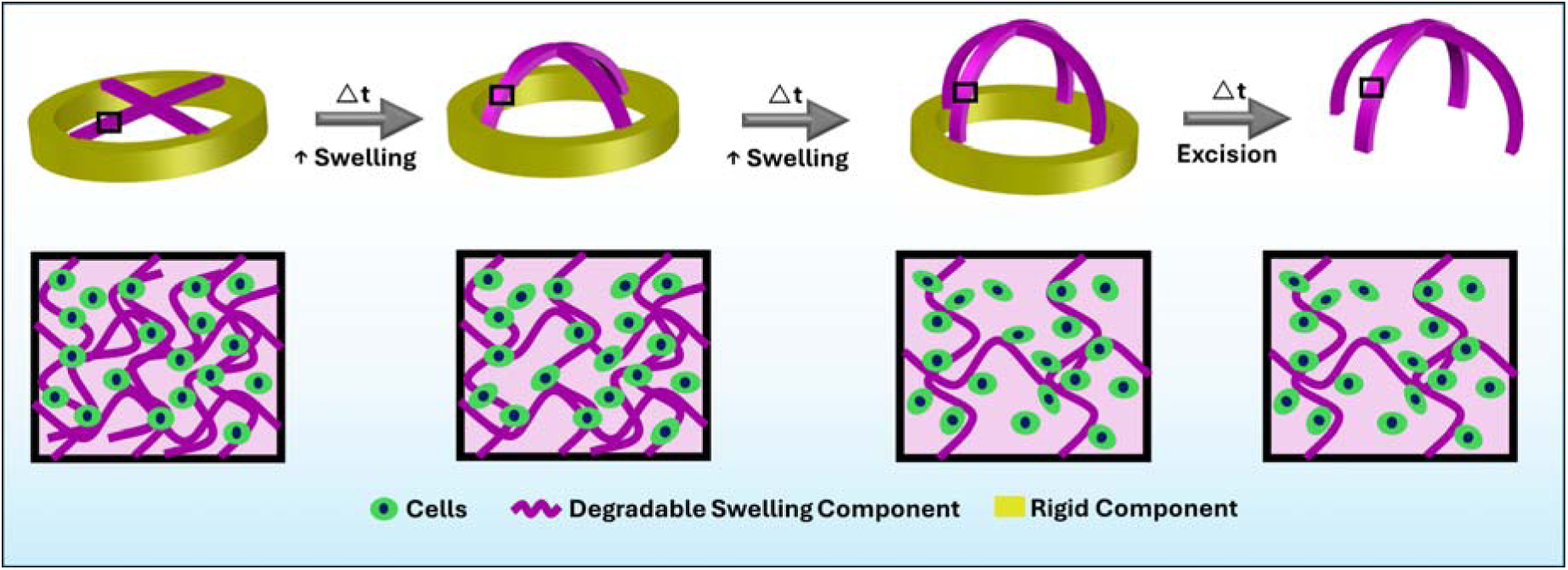
Conceptual illustration of the bio-Kirigami system. The bio-Kirigami construct comprises a rigid outer support ring (yellow) and an inner deformable, cell-laden component (magenta). Upon culture in medium, the inner component undergoes continuous shape morphing accompanied by tissue differentiation over time, ultimately forming complex tissue curvatures.

## References

[1] J.A. Espina, M.H. Cordeiro, E.H. Barriga, Tissue interplay during morphogenesis, Semin Cell Dev Biol 147 (2023) 12–23.

[2] M. Tozluoglu, Y. Mao, On folding morphogenesis, a mechanical problem, Philos Trans R Soc Lond B Biol Sci 375(1809) (2020) 20190564.

[3] S.B. Lemke, C.M. Nelson, Dynamic changes in epithelial cell packing during tissue morphogenesis, Current Biology 31(18) (2021) R1098–R1110.

[4] C.-P. Heisenberg, Y. Bellaïche, Forces in Tissue Morphogenesis and Patterning, Cell 153(5) (2013) 948–962.

[5] Y. Zhu, S. Deng, X. Zhao, G. Xia, R. Zhao, H.F. Chan, Deciphering and engineering tissue folding: A mechanical perspective, Acta Biomater 134 (2021) 32–42.

[6] A.J. Hughes, H. Miyazaki, M.C. Coyle, J. Zhang, M.T. Laurie, D. Chu, Z. Vavrušová, R.A. Schneider, O.D. Klein, Z.J. Gartner, Engineered Tissue Folding by Mechanical Compaction of the Mesenchyme, Developmental Cell 44(2) (2018) 165–178.e6.

[7] D. Wu, K.M. Yamada, S. Wang, Tissue Morphogenesis Through Dynamic Cell and Matrix Interactions, Annu Rev Cell Dev Biol 39 (2023) 123–144.

[8] I. Martyn, Z.J. Gartner, Expanding the boundaries of synthetic development, Developmental Biology 474 (2021) 62–70.

[9] M.D. Diaz-de-la-Loza, B.M. Stramer, The extracellular matrix in tissue morphogenesis: No longer a backseat driver, Cells Dev 177 (2024) 203883.

[10] S.J.P. Callens, R.J.C. Uyttendaele, L.E. Fratila-Apachitei, A.A. Zadpoor, Substrate curvature as a cue to guide spatiotemporal cell and tissue organization, Biomaterials 232 (2020) 119739.

[11] C.P. Heisenberg, Y. Bellaiche, Forces in tissue morphogenesis and patterning, Cell 153(5) (2013) 948–62.

[12] B. Schamberger, R. Ziege, K. Anselme, M. Ben Amar, M. Bykowski, A.P.G. Castro, A. Cipitria, R.A. Coles, R. Dimova, M. Eder, S. Ehrig, L.M. Escudero, M.E. Evans, P.R. Fernandes, P. Fratzl, L. Geris, N. Gierlinger, E. Hannezo, A. Iglic, J.J.K. Kirkensgaard, P. Kollmannsberger, L. Kowalewska, N.A. Kurniawan, I. Papantoniou, L. Pieuchot, T.H.V. Pires, L.D. Renner, A.O. Sageman-Furnas, G.E. Schroder-Turk, A. Sengupta, V.R. Sharma, A. Tagua, C. Tomba, X. Trepat, S.L. Waters, E.F. Yeo, A. Roschger, C.M. Bidan, J.W.C. Dunlop, Curvature in Biological Systems: Its Quantification, Emergence, and Implications across the Scales, Adv Mater 35(13) (2023) e2206110.

[13] C. Cui, D.-O. Kim, M.Y. Pack, B. Han, L. Han, Y. Sun, L.-H. Han, 4D printing of self-folding and cell-encapsulating 3D microstructures as scaffolds for tissue-engineering applications, Biofabrication 12(4) (2020) 045018.

[14] J. Malda, J. Visser, F.P. Melchels, T. Jungst, W.E. Hennink, W.J. Dhert, J. Groll, D.W. Hutmacher, 25th anniversary article: Engineering hydrogels for biofabrication, Adv Mater 25(36) (2013) 5011–28.

[15] A. Ding, O. Jeon, D. Cleveland, K.L. Gasvoda, D. Wells, S.J. Lee, E. Alsberg, Jammed Micro-Flake Hydrogel for Four-Dimensional Living Cell Bioprinting, Advanced Materials 34(15) (2022) 2109394.

[16] A. Roy, Z. Zhang, M.K. Eiken, A. Shi, A. Pena-Francesch, C. Loebel, Programmable Tissue Folding Patterns in Structured Hydrogels, Adv Mater (2023) e2300017.

[17] J. Lai, Y. Liu, G. Lu, P. Yung, X. Wang, R.S. Tuan, Z.A. Li, 4D bioprinting of programmed dynamic tissues, Bioactive Materials 37 (2024) 348–377.

[18] A. Ding, F. Tang, E. Alsberg, 4D Printing: A Comprehensive Review of Technologies, Materials, Stimuli, Design, and Emerging Applications, Chem Rev 125(7) (2025) 3663–3771.

[19] A. Ding, F. Tang, E. Alsberg, The Emerging 4D Printing of Shape Memory Thermomorphs for Self Adaptative Biomedical.pdf>, Adv Funct Mater (2025).

[20] A. Ding, S.J. Lee, R. Tang, K.L. Gasvoda, F. He, E. Alsberg, 4D Cell-Condensate Bioprinting, Small 18(36) (2022) 2202196.

[21] L. Ionov, 4D Biofabrication: Materials, Methods, and Applications, Adv Healthc Mater 7(17) (2018) e1800412.

[22] A. Ding, S.J. Lee, S. Ayyagari, R. Tang, C.T. Huynh, E. Alsberg, 4D biofabrication via instantly generated graded hydrogel scaffolds, Bioactive Materials 7 (2022) 324–332.

[23] A. Kirillova, R. Maxson, G. Stoychev, C.T. Gomillion, L. Ionov, 4D Biofabrication Using Shape Morphing Hydrogels, Advanced Materials 29(46) (2017).

[24] T. Pelluau, T. Brossier, M. Habib, S. Sene, G. Félix, J. Larionova, S. Blanquer, Y. Guari, 4D Printing Nanocomposite Hydrogel Based on PNIPAM and Prussian Blue Nanoparticles Using Stereolithography, Macromolecular Materials and Engineering 309(3) (2023).

[25] X. Guo, X. Ni, J. Li, H. Zhang, F. Zhang, H. Yu, J. Wu, Y. Bai, H. Lei, Y. Huang, J.A. Rogers, Y. Zhang, Designing Mechanical Metamaterials with Kirigami-Inspired, Hierarchical Constructions for Giant Positive and Negative Thermal Expansion, Adv Mater 33(3) (2021) e2004919.

[26] T.S. Tran, R. Balu, S. Mettu, N. Roy Choudhury, N.K. Dutta, 4D Printing of Hydrogels: Innovation in Material Design and Emerging Smart Systems for Drug Delivery, Pharmaceuticals (Basel) 15(10) (2022).

[27] H. Cho, D.-N. Kim, Controlling the stiffness of bistable kirigami surfaces via spatially varying hinges, Materials & Design 231 (2023).

[28] R. Sajjad, S.T. Chauhdary, M.T. Anwar, A. Zahid, A.A. Khosa, M. Imran, M.H. Sajjad, A review of 4D printing – Technologies, shape shifting, smart polymer based materials, and biomedical applications, Advanced Industrial and Engineering Polymer Research 7(1) (2024) 20–36.

[29] X.P. Hao, Z. Xu, C.Y. Li, W. Hong, Q. Zheng, Z.L. Wu, Kirigami-Design-Enabled Hydrogel Multimorphs with Application as a Multistate Switch, Adv Mater 32(22) (2020) e2000781.

[30] A. Rafsanjani, K. Bertoldi, Buckling-Induced Kirigami, Phys Rev Lett 118(8) (2017) 084301.

[31] M.A. Dias, M.P. McCarron, D. Rayneau-Kirkhope, P.Z. Hanakata, D.K. Campbell, H.S. Park, D.P. Holmes, Kirigami actuators, Soft Matter 13(48) (2017) 9087–9092.

[32] A.K. Brooks, S. Chakravarty, M. Ali, V.K. Yadavalli, Kirigami-Inspired Biodesign for Applications in Healthcare, Adv Mater 34(18) (2022) e2109550.

[33] L. Jin, A.E. Forte, B. Deng, A. Rafsanjani, K. Bertoldi, Kirigami-Inspired Inflatables with Programmable Shapes, Adv Mater 32(33) (2020) e2001863.

[34] Y. Zheng, I. Niloy, I. Tobasco, P. Celli, P. Plucinsky, Modelling planar kirigami metamaterials as generalized elastic continua, Proceedings of the Royal Society A: Mathematical, Physical and Engineering Sciences 479(2272) (2023).

[35] S. Pradhan, L. Ventura, F. Agostinacchio, M. Xu, E. Barbieri, A. Motta, N.M. Pugno, V.K. Yadavalli, Biofunctional Silk Kirigami With Engineered Properties, ACS Appl Mater Interfaces 12(11) (2020) 12436–12444.

[36] H. Choi, Y. Luo, G. Olson, P. Won, J.H. Shin, J. Ok, Y.J. Yang, T.i. Kim, C. Majidi, Highly Stretchable and Strain Insensitive Liquid Metal based Elastic Kirigami Electrodes (LM eKE), Advanced Functional Materials 33(30) (2023).

[37] A.J. Hughes, H. Miyazaki, M.C. Coyle, J. Zhang, M.T. Laurie, D. Chu, Z. Vavrusova, R.A. Schneider, O.D. Klein, Z.J. Gartner, Engineered Tissue Folding by Mechanical Compaction of the Mesenchyme, Dev Cell 44(2) (2018) 165–178 e6.

[38] Z.J. Wang, C.N. Zhu, W. Hong, Z.L. Wu, Q. Zheng, Cooperative deformations of periodically patterned hydrogels, Science Advances 3(9) (2017).

[39] S. Chen, J. Chen, X. Zhang, Z.Y. Li, J. Li, Kirigami/origami: unfolding the new regime of advanced 3D microfabrication/nanofabrication with "folding", Light Sci Appl 9 (2020) 75.

[40] H. Li, W. Zhang, X. Liao, L. Xu, Kirigami enabled reconfigurable three-dimensional evaporator arrays for dynamic solar tracking and high efficiency desalination, Sci Adv 10(26) (2024) eado1019.

[41] Y. Tang, Y. Li, Y. Hong, S. Yang, J. Yin, Programmable active kirigami metasheets with more freedom of actuation, Proc Natl Acad Sci U S A 116(52) (2019) 26407–26413.

[42] A.M. Abdullah, X. Li, P.V. Braun, J.A. Rogers, K.J. Hsia, Kirigami Inspired Self Assembly of 3D Structures, Advanced Functional Materials 30(14) (2020).

[43] C. Dai, Y. Rho, K. Pham, B. McCormick, B.W. Blankenship, W. Zhao, Z. Zhang, S.M. Gilbert, M.F. Crommie, F. Wang, C.P. Grigoropoulos, A. Zettl, Kirigami Engineering of Suspended Graphene Transducers, Nano Letters 22(13) (2022) 5301–5306.

[44] P. Li, X. Dou, C. Feng, H. Schönherr, Enhanced cell adhesion on a bio-inspired hierarchically structured polyester modified with gelatin-methacrylate, Biomaterials Science 6(4) (2018) 785–792.

[45] R. Zhao, S. Lin, H. Yuk, X. Zhao, Kirigami enhances film adhesion, Soft Matter 14(13) (2018) 2515–2525.

[46] K. Wei, A. Roy, S. Ejike, M.K. Eiken, E.M. Plaster, A. Shi, M. Shtein, C. Loebel, Magnetoactive, Kirigami-Inspired Hammocks to Probe Lung Epithelial Cell Function, Cell Mol Bioeng 17(5) (2024) 317–327.

[47] E.E. Evke, D. Meli, M. Shtein, Developable Rotationally Symmetric Kirigami Based Structures as Sensor Platforms, Advanced Materials Technologies 4(12) (2019).

[48] L.J. Macdougall, M.M. Perez-Madrigal, J.E. Shaw, M. Inam, J.A. Hoyland, R. O’Reilly, S.M. Richardson, A.P. Dove, Self-healing, stretchable and robust interpenetrating network hydrogels, Biomater Sci 6(11) (2018) 2932–2937.

[49] L.D. Solorio, C.D. Dhami, P.N. Dang, E.L. Vieregge, E. Alsberg, Spatiotemporal Regulation of Chondrogenic Differentiation with Controlled Delivery of Transforming Growth Factor-β1 from Gelatin Microspheres in Mesenchymal Stem Cell Aggregates, STEM CELLS Translational Medicine 1(8) (2012) 632–639.

[50] S. Chen, Z. Liu, H. Du, C. Tang, C.Y. Ji, B. Quan, R. Pan, L. Yang, X. Li, C. Gu, X. Zhang, Y. Yao, J. Li, N.X. Fang, J. Li, Electromechanically reconfigurable optical nano-kirigami, Nat Commun 12(1) (2021) 1299.

[51] Z. Zou, B. Zhang, X. Nie, Y. Cheng, Z. Hu, M. Liao, S. Li, A sodium alginate-based sustained-release IPN hydrogel and its applications, RSC Adv 10(65) (2020) 39722–39730.

[52] H. Omidian, S.D. Chowdhury, R.L. Wilson, Advancements and Challenges in Hydrogel Engineering for Regenerative Medicine, Gels 10(4) (2024).

[53] X. Guan, M. Avci-Adali, E. Alarcin, H. Cheng, S.S. Kashaf, Y. Li, A. Chawla, H.L. Jang, A. Khademhosseini, Development of hydrogels for regenerative engineering, Biotechnol J 12(5) (2017).

[54] X. Bai, M. Gao, S. Syed, J. Zhuang, X. Xu, X.Q. Zhang, Bioactive hydrogels for bone regeneration, Bioact Mater 3(4) (2018) 401–417.

[55] A. Ding, K.L. Gasvoda, D.S. Cleveland, E. Alsberg, Cell Contractile Force-Mediated Morphodynamical Tissue Engineering via 4D Printed Degradable Hydrogel Scaffolds, bioRxiv (2025).

[56] C.T. Huynh, Z. Zheng, M.K. Nguyen, A. McMillan, G. Yesilbag Tonga, V.M. Rotello, E. Alsberg, Cytocompatible Catalyst-Free Photodegradable Hydrogels for Light-Mediated RNA Release To Induce hMSC Osteogenesis, ACS Biomaterials Science & Engineering 3(9) (2017) 2011–2023.

[57] J.E. Samorezov, E.B. Headley, C.R. Everett, E. Alsberg, Sustained presentation of BMP-2 enhances osteogenic differentiation of human adipose-derived stem cells in gelatin hydrogels, Journal of Biomedical Materials Research Part A 104(6) (2016) 1387–1397.

[58] O. Jeon, D.S. Alt, S.M. Ahmed, E. Alsberg, The effect of oxidation on the degradation of photocrosslinkable alginate hydrogels, Biomaterials 33(13) (2012) 3503–14.

[59] O. Jeon, Y.B. Lee, S.J. Lee, N. Guliyeva, J. Lee, E. Alsberg, Stem cell-laden hydrogel bioink for generation of high resolution and fidelity engineered tissues with complex geometries, Bioact Mater 15 (2022) 185–193.

[60] M.K. Nguyen, O. Jeon, M.D. Krebs, D. Schapira, E. Alsberg, Sustained localized presentation of RNA interfering molecules from in situ forming hydrogels to guide stem cell osteogenic differentiation, Biomaterials 35(24) (2014) 6278–6286.

[61] A. Ding, S.J. Lee, R. Tang, K.L. Gasvoda, F. He, E. Alsberg, 4D Cell-Condensate Bioprinting, Small 18(36) (2022) e2202196.

[62] L.D. Solorio, C.D. Dhami, P.N. Dang, E.L. Vieregge, E. Alsberg, Spatiotemporal regulation of chondrogenic differentiation with controlled delivery of transforming growth factor-beta1 from gelatin microspheres in mesenchymal stem cell aggregates, Stem Cells Transl Med 1(8) (2012) 632–9.

[63] A.K. Kudva, A.D. Dikina, F.P. Luyten, E. Alsberg, J. Patterson, Gelatin microspheres releasing transforming growth factor drive in vitro chondrogenesis of human periosteum derived cells in micromass culture, Acta Biomater 90 (2019) 287–299.

