## Supplementary Information for "A 4D Bio-Kirigami Strategy for Engineering Complex Tissue Curvatures"

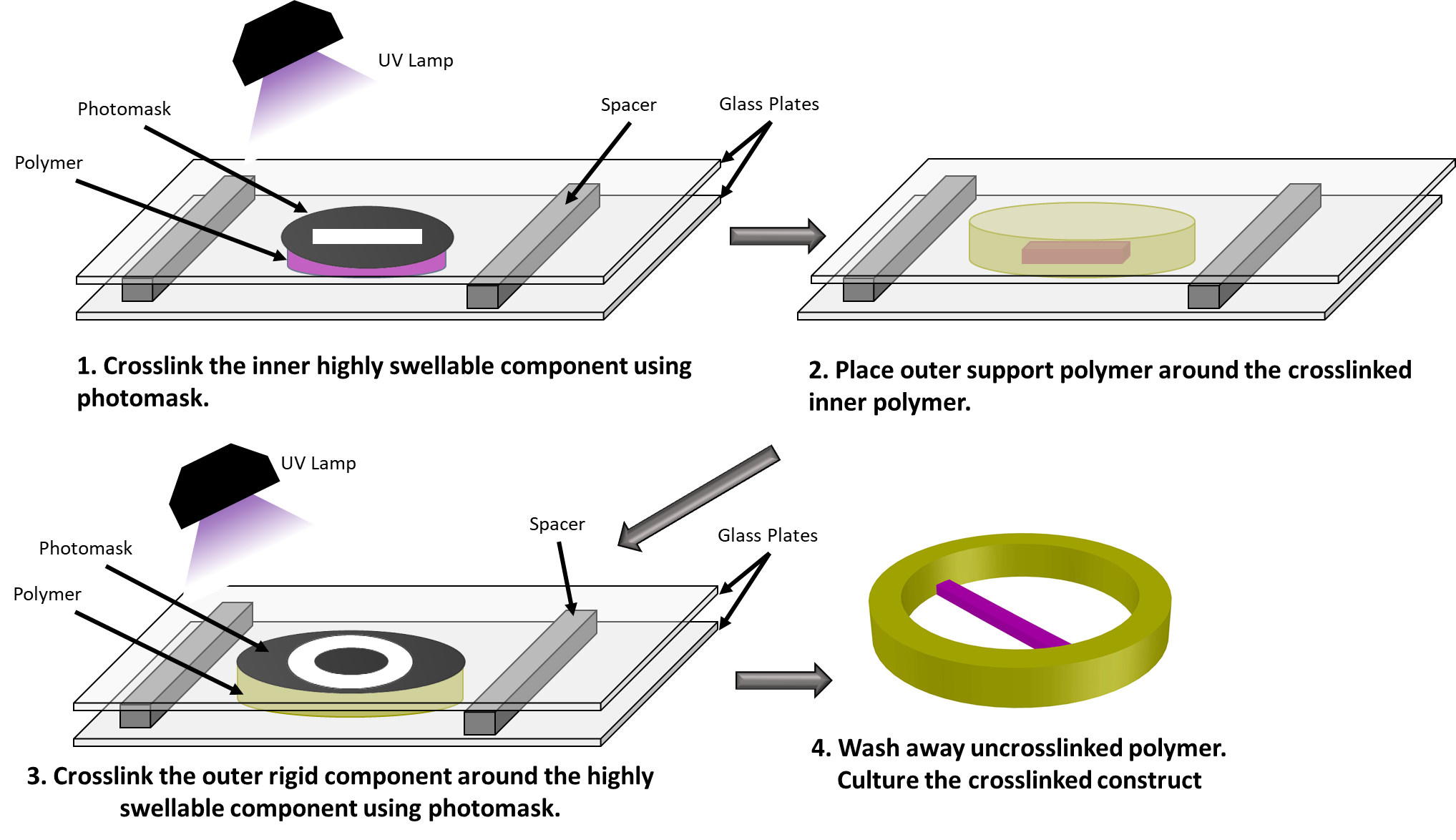

**Figure S1.** Schematic illustration of the fabrication process for bio-Kirigami constructs.

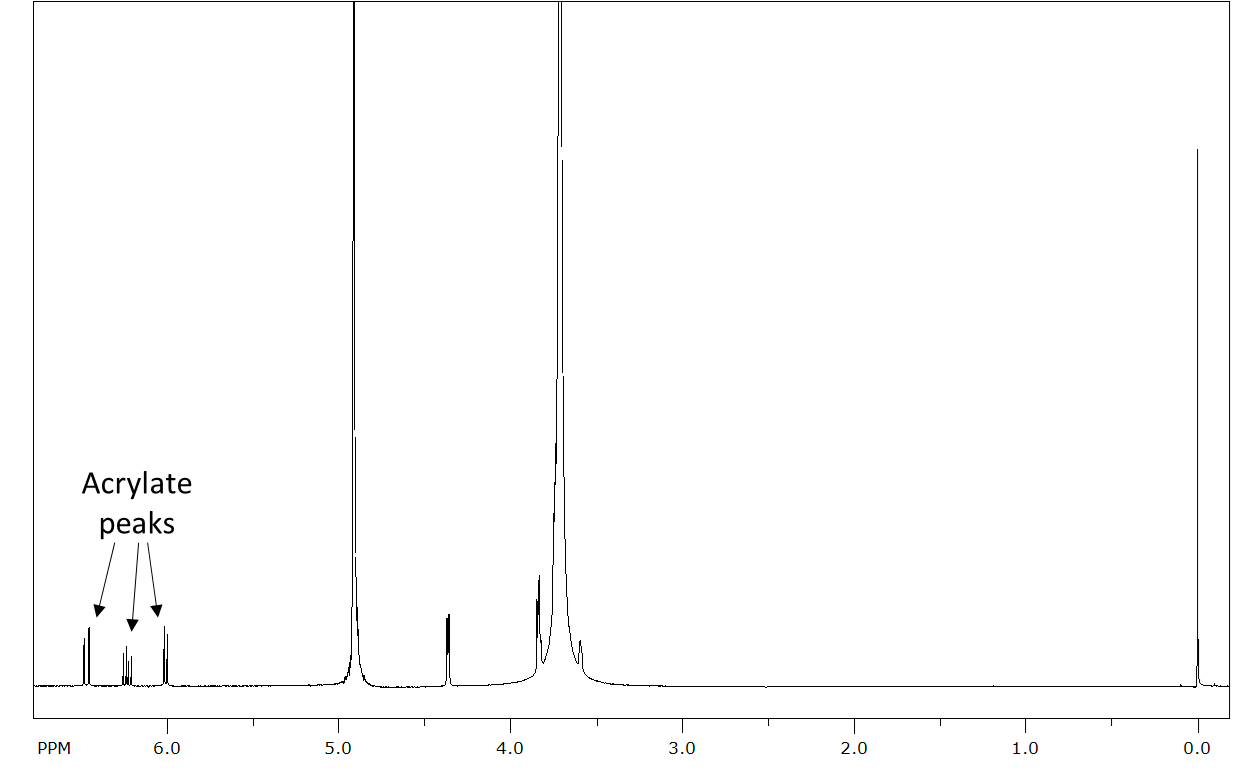

**Figure S2**. ^1^H NMR spectrum of PEGDA in D_2_O.

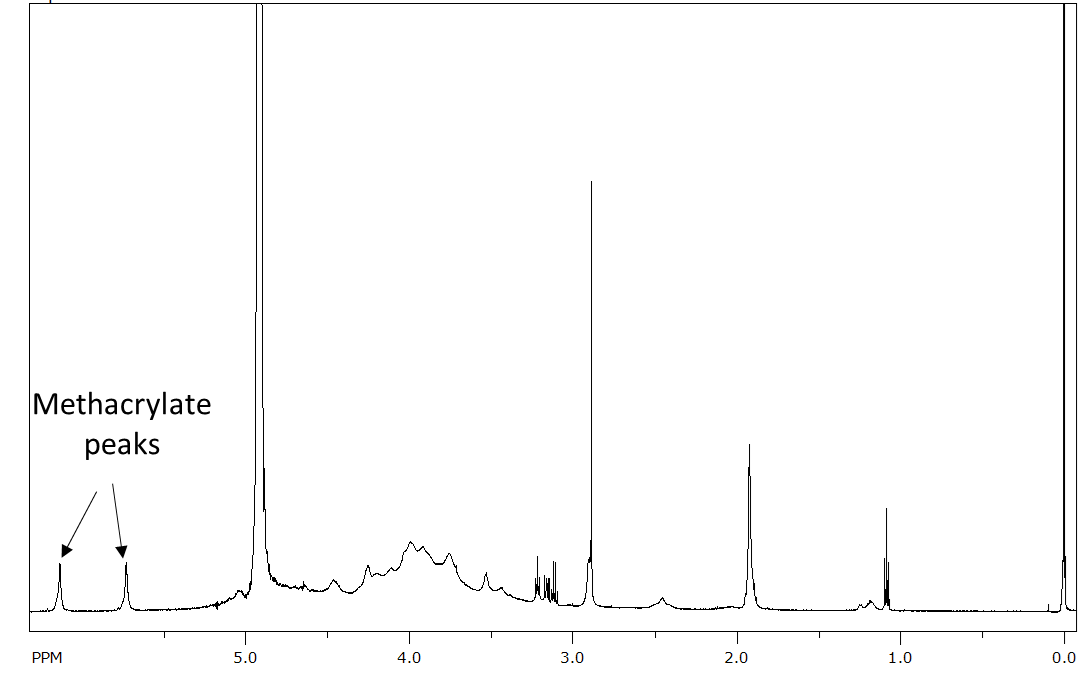

**Figure. S3.** ^1^H NMR spectrum of OMA in D_2_O.

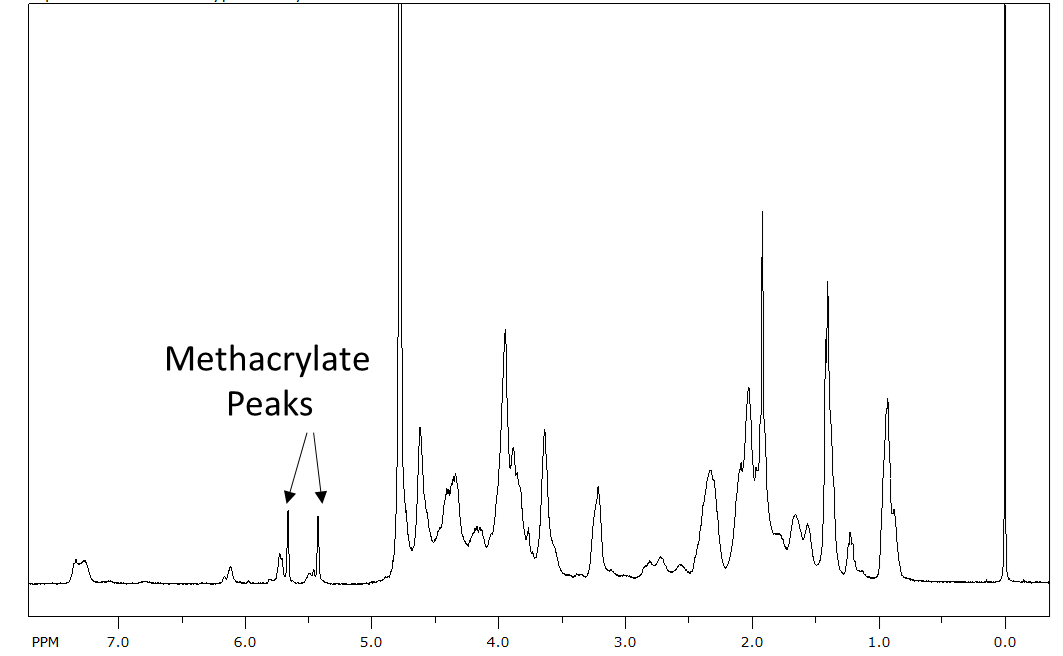

**Figure S4.** ^1^H NMR spectrum of GelMA in D_2_O.

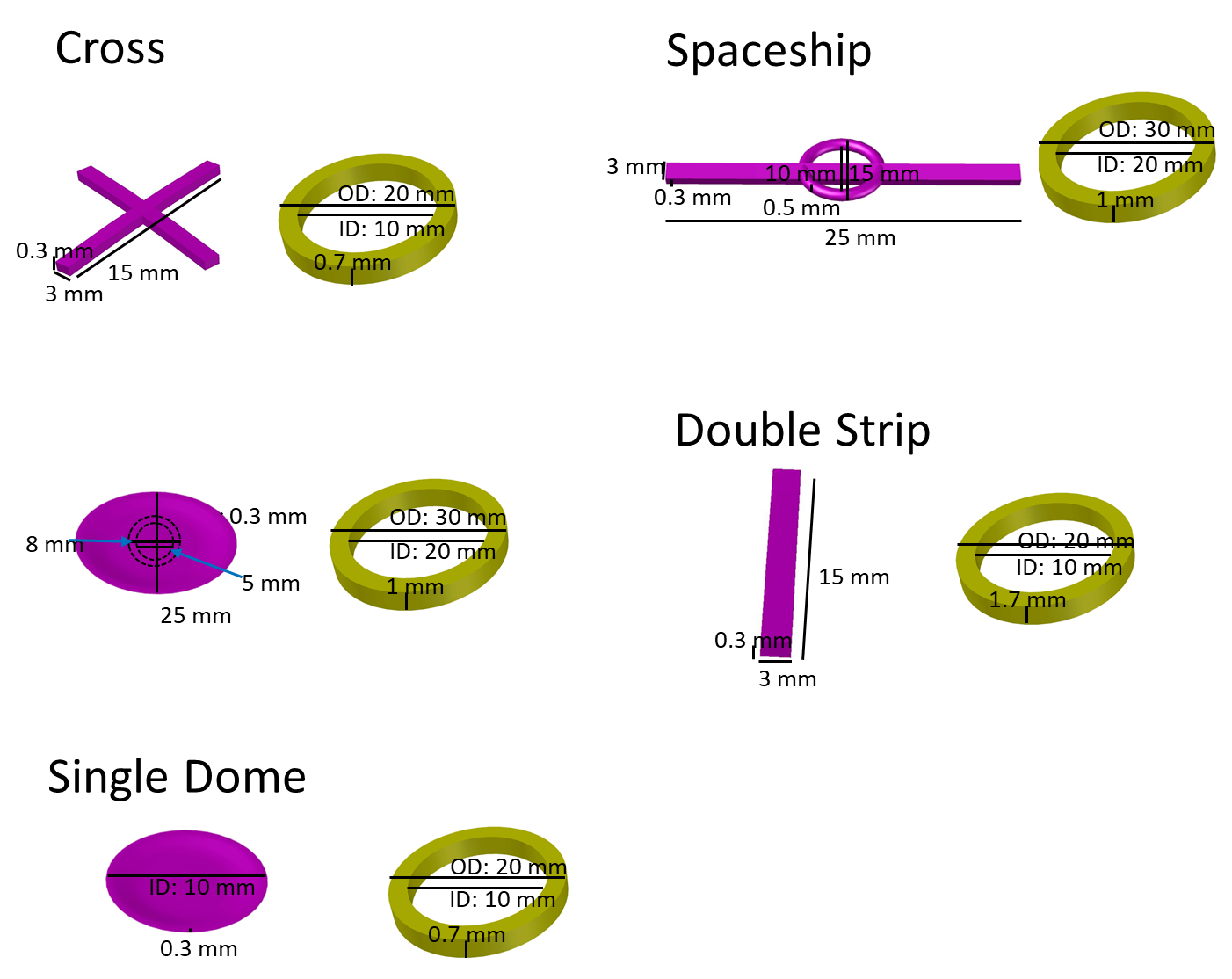

**Figure S5**. Dimensions of cell-free bio-Kirigami constructs.

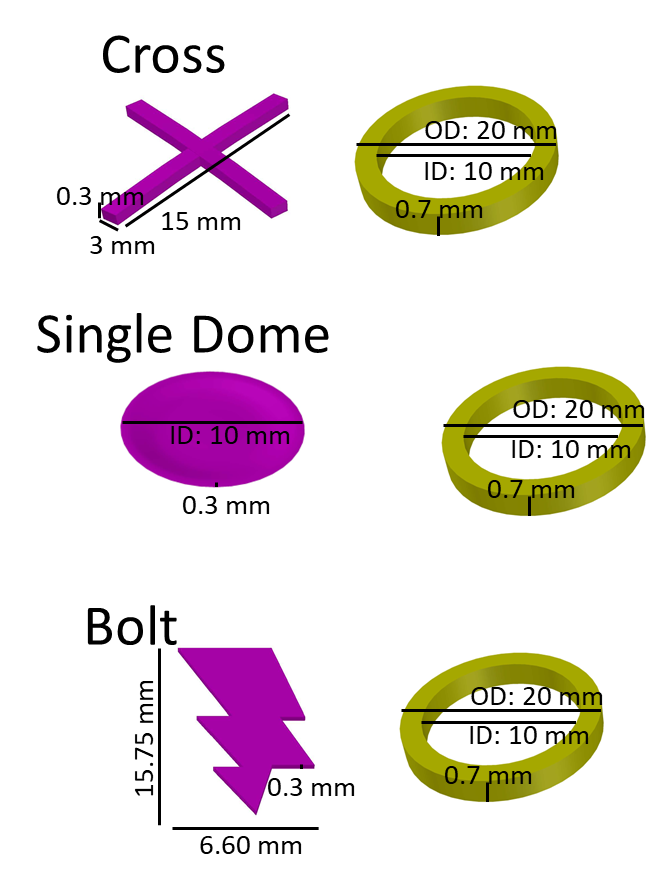

**Figure S6**. Dimensions of cell-laden bio-Kirigami constructs.
